# Short-term performance prediction in team sports using tracking data

**DOI:** 10.1101/2025.09.18.677041

**Authors:** Arnaud Odet, Thomas Bechard, Sebastien Déjean, Cristian Pasquaretta

**Author notes:** {, }, {, }.

## Abstract

This research addresses short-term performance prediction in Rugby Union using player tracking data. Rugby’s unique characteristics as a conquest sport with strict offside rules make the ball position a crucial performance indicator. We used event data to identify ball carriers in offensive phase and considered their locations as proxy for the ball. We then aim to predict this performance indicator. We formulated this challenge as a time series prediction problem where player positions (multivariate) predict performance (univariate). Our results suggest using player tracking data improves the prediction ability.

## Introduction

The prediction of performance in sport science has become increasingly important with the advancement of player tracking technologies and analytics (Kovalchik, 2022). Performance in sport can be defined and analyzed at different scales: (i) long-term (e.g. over a complete season, with studies about roster constitution Pérez-Toledano et al., 2019 or fatigue management Vanrenterghem et al., 2017), (ii) medium-term (e.g. over a game, with emphasis on feature importance Miljkovic et al., 2010, Odet et al., 2025), or short-term (e.g. over an action, Cervone et al., 2014). This study focuses on this third prediction task in the context of team sports, with a specific focus on Rugby Union, where the dynamic nature of play presents unique analytical opportunities and challenges. Our aim is to incorporate collective player behavior to predict a performance feature.

We formulate this objective as a time series prediction task, in which multivariate tracking data predicts univariate performance outcome. This formulation is particularly challenging in the context of team sports because the tracking data encode not only the temporal evolution of player movements but also the spatial interactions and collective coordination among them. Players’ movements in rugby are interconnected. Each player’s behavior is shaped by their role and by how they adapt to the movements of others, creating complex patterns that evolve over time. To properly capture these dependencies, models must go beyond basic statistical methods and represent both temporal dynamics and interactions between players. Recent studies have shown that deep learning models perform well in modeling time-dependent data, and that multivariate approaches can directly learn relationships between multiple movement series and time-varying interactions, outperforming traditional forecasting methods (Cao et al., 2020). These advances motivate our use of multivariate time series models to capture the effect of collective player movement on performance prediction in Rugby Union tracking data.

Rugby Union, as a conquest sport, is characterized by strict rules governing player positioning, particularly the offside rule (in short, a player is offside when positioned in front of the teammate who last played the ball and can not dispute it). These rules make ball position a crucial indicator of territorial advantage and play development. The relevance of ball position as a performance indicator in Rugby Union is strongly justified due to its direct influence on match outcomes and tactical strategies (Bennett et al., 2019; Colomer et al., 2020). In addition, linebreaks are identified as a crucial offensive indicator, as they directly contribute to advancing the ball’s position into advantageous opposition territory (Schoeman & Schall, 2019). Despite their importance, ball tracking data is often unavailable or incomplete in professional sports settings, creating a significant challenge for performance analysis.

To address this limitation, we combined manually annotated event data with player GPS tracking data and provide a method to estimate ball position in Rugby Union.

By leveraging these data in a predictive framework, it becomes possible to quantify aspects of team coordination and tactical decision-making, allowing for “what if” simulation analyses and continuous time play valuation. Such an approach have potential applications in both real-time game analysis and post-match performance evaluation, providing valuable tools for coaches, analysts, and sports scientists.

Contributions: we identified three main contributions of this work:

i. Short-term Performance Prediction: We formulated short-term performance prediction in Rugby Union as a multivariate to univariate time series forecasting task and demonstrated the relevance of using complex high-frequency tracking data.
ii. Publicly Available Dataset: We constructed our dataset using a novel preprocessing pipeline that includes a ball position estimation algorithm combining tracking and event data, addressing the challenge of incomplete ball tracking data in team sports.
iii. Benchmarking of several state-of-the-art time series models: We benchmark several state-of-the-art models, including an exploration of the hyperparameters space, versus popular vanilla models and naive baselines.

### Related Work

#### Tracking Data for Sports Performance Prediction

Initial applications of tracking data in team sports primarily focused on quantifying physical performance and conducting time-motion analysis (Ferraz et al., 2023). This field has since undergone a paradigmatic shift, moving towards using high-resolution spatio-temporal data for more sophisticated tactical analysis and understanding collective movement patterns among players (Low et al., 2020; Perše et al., 2009). A key area of research involves deriving performance indicators from spatial and temporal aggregations of tracking data, utilizing metrics such as team centroid, space control, and other domain-specific features (Clemente et al., 2013; Low et al., 2020). Building upon this, a significant development is the concept of continuous-time within-play valuation. This includes the development of frameworks for estimating Expected Possession Value (EPV), which quantifies the expected number of points an offense will score on a possession conditional on its evolution, using a multiresolution stochastic process model for basketball (Cervone et al., 2014). The EPV concept has been adapted to soccer with a decoupled deep learning approach that provides a frame-by-frame estimation of the expected outcome of a possession (Fernández, 2020). Similarly, models for estimating Expected Points (EP) and Win Probability (WP) continuously throughout a play have been developed for American football, using Long Short-Term Memory (LSTM) networks to model outcomes based on player locations and trajectories (Yurko et al., 2019). Machine learning techniques, including Graph Neural Networks (GNNs) and Recurrent Neural Networks (RNNs), allows to automate the identification of tactical patterns and summarize complex player interactions, reducing the reliance on manual annotation (Anzer et al., 2022; Tian et al., 2019). These advanced data-driven models enable the rating of specific actions not just by their outcome probability, but more comprehensively by quantifying the value they add or subtract from the expected outcome of a possession or play (Dick et al., 2021).

While machine learning has been increasingly applied to performance prediction in sports, the specific task of trajectory prediction, i.e. estimating the future position of a ball or player, remains relatively underexplored in the literature. This task is particularly important in team sports like rugby, where anticipating the movement of the ball and players is critical for strategic planning, performance analysis, and real-time decision making. Trajectory prediction has been successfully applied in other domains such as robotics, autonomous vehicles, and aerospace, where forecasting object motion is essential for navigation and control (references). Inspired by these approaches, we developed a machine learning model that predicts the trajectory of the rugby ball using both player and ball coordinates, enabling the estimation of ball position several seconds into the future. This approach captures the complex interactions on the field and provides a data-driven tool to support tactical analysis and game understanding. The present work aims at leveraging the sequential nature of tracking data for modeling short-term performance in Rugby Union and brings novelty by benchmarking state-of-the-art time series models on this task.

#### Time series Forecasting

Time series analysis involves studying collections of observations ordered by time, with applications across various fields including energy, economics, and meteorology. The main challenge is to understand the temporal dynamics in time series, which includes decomposition, seasonality, long-term dependencies, etc. It has been a very dynamic, long-standing field of study (Klein, 1997), with traditional statistical methods (ARMA and descendants, see Box et al., 1978) and advances in Deep Learning with RNN (Rumelhart et al., 1986), LSTM (Hochreiter & Schmidhuber, 1997), Transformer (Vaswani et al., 2017) and more recently Mamba (Gu & Dao, 2023). Transformer models have been adapted for Time series analysis for their ability to capture long-range dependencies, with a wide focus on the attention mechanism (see review by Wang et al. (2024)). PatchTST (Nie et al., 2023) improves long-term forecasting accuracy by segmenting time series into subseries-level patches. Non-stationary Transformers (Y Liu et al., 2022) uses methods like De-stationary Attention to handle non-stationarity in time series, outperforming stationarization techniques. Pyraformer (S Liu et al., 2022) employs a multi-resolution pyramidal attention structure to efficiently model dependencies across various time scales. Crossformer (Zhang & Yan, 2023) explicitly addresses both cross-time and crossdimension dependencies via a dimension-segment-wise embedding and two-stage attention layer. In contrast to temporal token-based methods, iTransformer (Y Liu et al., 2023) proposes an inverted architecture that applies attention on variate tokens to capture multivariate correlations and uses feed-forward networks for learning variate-centric representations. Zeng et al. (2022) questioned the necessity of using Transformers for long-term forecasting, suggesting that simpler linear models can also achieve competitive results. These models are compared on reference datasets (energy, foreign exchange, weather, etc. see Wang et al., 2024) The present study applies these models to sequences expected to present distinct temporal dynamics than those of the reference datasets.

#### Ball position estimation

While tacking ball position from videos is a flourishing topic in computer vision (Ren et al., 2009), effort to estimate the ball trajectory without optical or tracking data remain limited. Amirli and Alemdar (2022) used neural network on Soccer players’ tracking data and reached a MAE greater than 5 meters. Kim et al. (2023) combined set transformers and hierarchical architecture to achieve a 3.6 meters MAE. Our work differs as we used heuristics leveraging event data.

### Methodology

#### Problem Formulation

Let *X*_*i,j*_ be the feature vector of player *j* ∈ {1,…, 15} at time 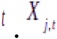. is 4-dimensional, as it contains the position ({*x*_*j*_}_*t*_ and {*y*_*j*_}_*t*_ coordinates) and the velocity 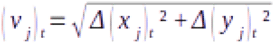 where 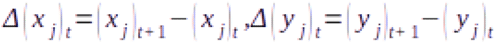 and the goal angle *ϕ*_j_ = arctan (*Δ*(*y*_*j*_) / *Δ* (*x*_*j*_)) of Player *j* (the goal angle is the angle between the actual trajectory and a trajectory consisting in running straight to the opponent try line, i.e. with *Δ* {*x*_*j*_} > 0 and *Δ* {*y*_*j*_} = 0). We note *s* the sequence length, *p* the prediction length (both in tenth of second), and *P*_*t*_ the relative value of our performance feature at time *t*. It is obtained by subtracting the value of the performance feature at the end of the known sequence (time *t*_0_ + *s* for any sequence starting at time *t*_0_) to the value of the performance feature at any time t ∈ [*t*_0_, *t*_0_, + *s* + *p*], materializing the progress of the action. Here, we chose the ball coordinate over the *x* axis as performance feature, as it reflects the territorial advantage. Let *S*_*t*_ = *P* _*t* + *s* + *p*_ be the performance variable associated with the sequence [*Xt*,…, *Xt* + *s*], where *X*_*t*_ = {*X*_[1,*t*]_,…, *X*_[15,*t*]_,*P*_*t*_} is the feature vector at time *t* (*X*_*t*_ has 61 features)

Our objective can be formulated as predicting *S*_*t*_ given {*X*_*t*_,…, *X*_*t*+ *s*_}. Alternatively, it can be reformulated as: Predicting {*P*_*t*+ *s* + 1_,…, _*t*+ *s* + *p*_} given {*X*_*t*_,…, *X*_*t*+ *s*_} (multivariate to univariate time series prediction task), where methods are compared using the Final Displacement Error (FDE) metric, as the FDE only considers the last point of a trajectory.

Formally, the FDE is defined as 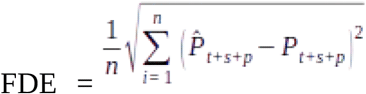 where 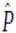 is the estimated performance and *t*_*i*_ indicates the start of sequence *i* ∈ {1,…,*n*}.

#### Dataset construction

We used a dataset containing 47 French Pro D2 games of a given team. A distribution of these games by location and outcome can be found in Table 1.

**Table 1:** Games locations and outcomes.

|  | Win | Draw | Loss | **Total** |
| --- | --- | --- | --- | --- |
| Away | 4 | - | 13 | **17** |
| Home | 24 | 2 | 4 | **30** |
| **Total** | **28** | **2** | **17** | **47** |

For these games, we dispose of (i) event data and (ii) tracking data. Event data are manually labeled by the team’s video analysts. Each entry corresponds to an event, and comprises its type (pass, tackle, kick, substitution, yellow or red card, etc.), timestamp, and the identity of the player performing it. Tracking data consists of the GPS position of all 23 players (including 8 substitutes) during the game, with a frequency of 10 Hz. This data is only available for the team which provided it. We have no data about the opposing team.

From event data, we identified the specific time intervals at which an action starts and finishes. We gathered 22 hours of playtime (4542 actions), from which we only considered offensive situations (11 hours - 2248 actions) as our ball position estimation method can only be applied when the analyzed team is in possession of the ball. We also used event data to remove from our analysis sequences in which the team is in numerical inferiority (due to yellow or red card) for the sake of simplicity, we set aside sequences with missing or aberrant values and actions either shorter than 3.2 seconds (our shortest time horizon, see below) or without identified ball handler (required to quantify performance, see below). After this cleaning step, 8 hours (1332 actions) of offensive play remain available for analysis.

We then split our data into sequences of different time horizons, to verify the robustness of findings. A time horizon consists in sequence length (the number of historical time steps used as input for prediction) and prediction length (the number of following time steps to be forecasted). Sequences are then split into train (70%), validation (15%) and test (15%) sets. Table 2 displays the number of actions and sequences for each time horizon.

**Table 2:** Number of actions and sequences per time horizon.

| Time Horizon | Actions | Sequences |
| --- | --- | --- |
| 16-16 | 1,332 | 112,325 |
| 32-32 | 1,093 | 90,033 |
| 64-64 | 735 | 57,629 |
| 128-128 | 303 | 23,771 |

We transformed tracking Latitude and Longitude to x and y coordinates where the (0, 0) coordinate is the lower-left corner from top view of the pitch. We rotate second halves, so the analyzed team always attacks from left to right (increasing x for successful attacks), and we computed the velocity and goal angle of all 15 players playing the sequence (identified based on the starting lineup and the changes registered in event data).

##### Ball position extraction method

We used event data to estimate the position of the ball. For each possible type of events (approx. 30 are used by the technical staff), we identified if the event is a ball handler event (e.g. pass, breakthrough, etc.) or not (e.g. support, tackle, etc.). For ball handler events, we added a distinction between “give” event, where the ball handler loses control of the ball soon after the event is recorded (e.g. passes, kicks, etc.) and “keep” events, where the ball handler maintain control of the ball (e.g. breakthrough, contacts, etc.). This event classification was obtained from the technical staff.

We then used for each ball handler event the coordinates of the ball handler as coordinates of the ball for the timestamp of the corresponding event. For all timestamps between two ball handler events (meaning the vast majority of timestamps, as ball handler events do not occur at the same frequency the tracking data is captured), we used the coordinates of the ball handler of the first (respectively second) event if the event is considered a “keep” (respectively “give”) event, and applied Gaussian filter to smooth ball trajectories.

To the best of our knowledge, this dataset is the only publicly available dataset containing Rugby Union Tracking data. We believe its size (47 games) and frequency (10Hz) make it valuable for the broader scope of team sports performance analysis.

### Experiments

#### Experimental Setup

We compared the following methods: two naive baselines (Stand Still: 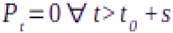 where *t*_0_ is the start of the sequence, Constant Velocity: 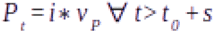 where 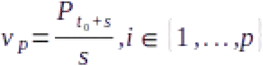 - an illustration of target series and baselines is displayed in Figure 1), 2 vanilla models (LSTM Hochreiter and Schmidhuber, 1997 and Transformer Vaswani et al., 2017) and several state-of-the-art time series models, highlighted in Wang et al. (2024): iTransformer (Y Liu et al., 2023), PatchTST (Nie et al., 2023), Non-Stationary Transformer (Y Liu et al., 2022), Crossformer (Zhang & Yan, 2023), Pyraformer (S Liu et al., 2022) and DLinear (Zeng et al., 2022). For the time series and vanilla models, we ran the two experiments: the first where only our prediction variable is used to predict itself (univariate to univariate prediction task) and the second where the whole feature vector is used to predict the target variable (multivariate to univariate prediction task). The comparison between both experiments aims at quantifying the prediction ability gained by using tracking data.

**Fig. 1:**
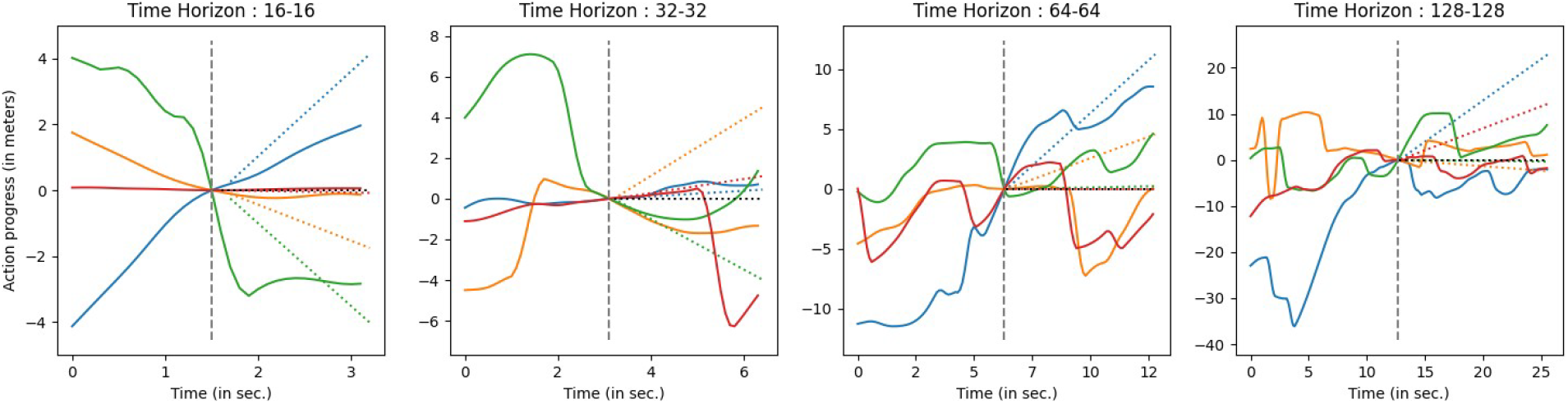
Illustration of 4 randomly selected target series P over different time horizons. The vertical dashed gray line represents the separation between known sequence on the left and prediction sequence on the right, the horizontal dotted black line represents the Stand Still baseline and the colored dotted lines the Constant Velocity baseline associated with the serie displayed with the same color.

#### Training procedure

Due to the lack of prior knowledge about which model would best fit our data, the considerable size of the hyperparameters space, and for a reasonable use of computing resources, we designed a halving / pruning protocol (Jamieson & Talwalkar, 2015). For each model, we conducted various experiments based on common hyperparameters ranges, taking into consideration authors’ suggestions. We limited the training budget to 100 epochs with an early stopping criterion set at 8, and every 4 epochs, we compared the validation loss of all “alive” models and pruned half of them. We used *adamw* optimizer (Loshchilov & Hutter, 2017), MSE loss, and a customized LR scheduler with warm-up and restarts (applying a decay of 0.5 at each restart) aligned with our pruning cycle.

#### Testing procedure

At the end of training, we evaluated over the test set the models of which training was terminated either by early stopping or by reaching the training budget (either case meaning the experiment completed the training). We found that in rare cases, during training, a given model *M* having reached a validation loss earlier *z*_0_ during its training arrives at pruning step with a validation loss *z*_0_ > *z*_*P*_ > *z*_0_, where *z*_*P*_ is the pruning threshold. *M* is therefore pruned, and budget is allocated to other models, of which some will never reach *z*_0_ and yet be tested. In such cases, we reintegrated *M* for testing.

## Results and Discussions

### Training and pruning

Figure 2 (and Appendix 1) displays, for each time horizon, the best epoch and the MSE on the validation set for the 5 best experiments by model, for univariate and multivariate inputs. For most models, the best epoch is earlier for multivariate inputs than for univariate inputs. This observation is not surprising as considering more features to predict the same outputs can be expected to lead more easily to overfitting. This finding gets more visible with longer time horizons, which can be explained by the lower number of sequences (see Table 2). The consequence of the best epoch being reached earlier is fewer multivariate inputs models will be pruned, and therefore more will reach testing. During training, in most of the cases, univariate inputs models reached lower MSE on the validation set than their multivariate counterparts, which could indicate a better ability to fit the data. However, for shorter time horizons (left part of Figure 2), multivariate models reached lower MSE, with Crossformer (Zhang & Yan, 2023) for both univariate and multivariate reaching the lowest MSE. For longer time horizons (right part of Figure 2), Pyraformer (S Liu et al., 2022) for univariate and Crossformer (Zhang & Yan, 2023) and Patch TST (Nie et al., 2023) for multivariate performed best.

**Fig. 2:**
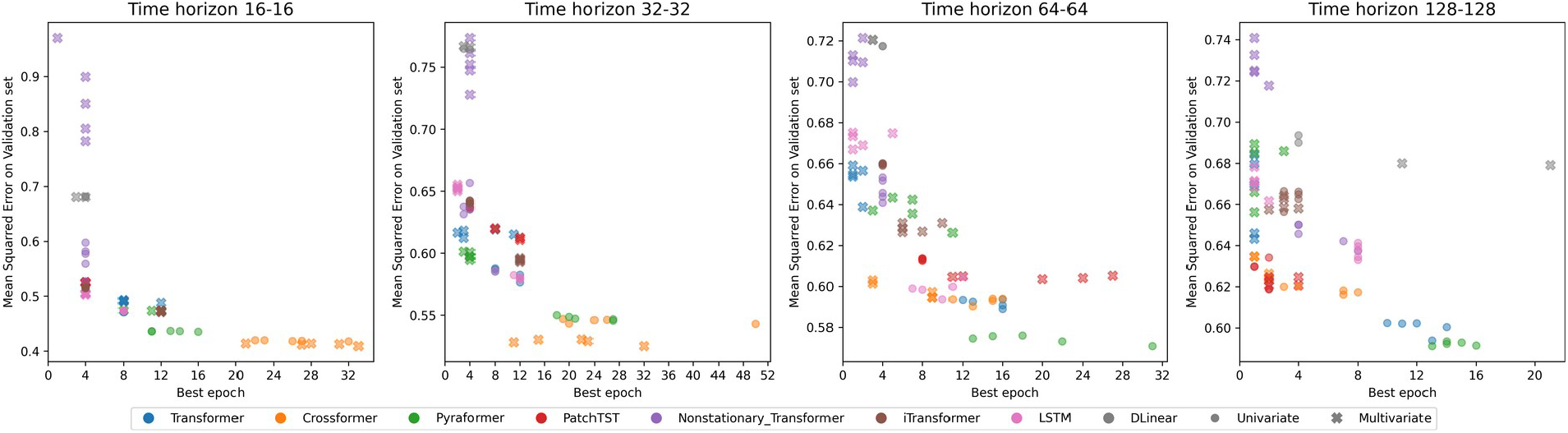
MSE and best epoch of the 5 best experiments for each model over the validation set.

### Testing

Figure 3 (and Appendix 2) displays, for each time horizon, the distribution of relative improvements in FDE over the test set against the best performing baseline for models that reached the test step. Those models are those that either completed training or were reintegrated.

**Fig. 3:**
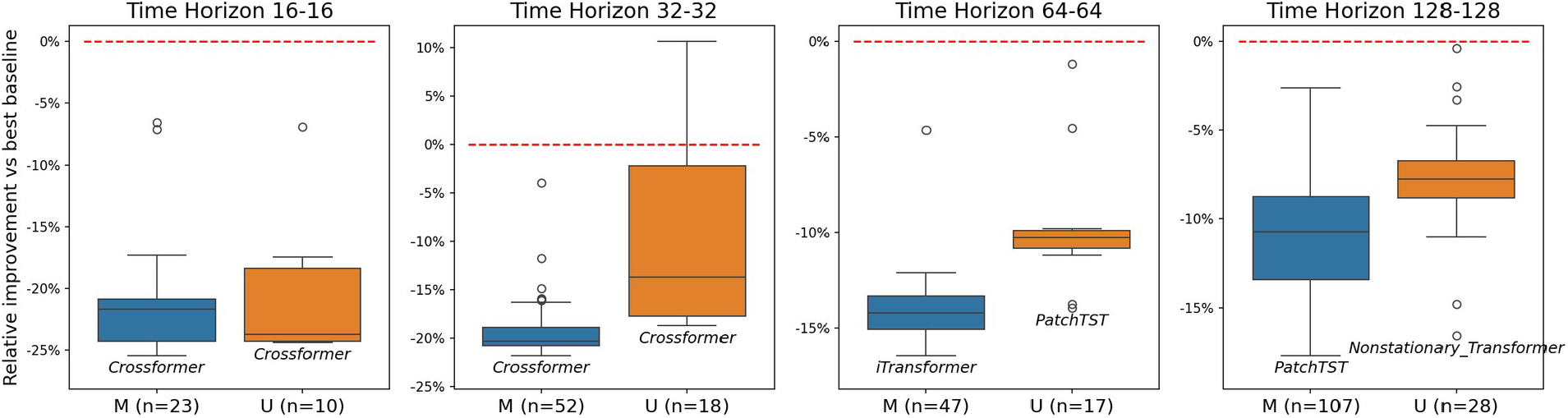
Distribution of FDE relatve improvement (in %) vs best baseline for Multivariate (M) and Univariate (U) inputs models for each time horizon. Each graph represents the results of the test procedure over the corresponding time horizon. The number in parenthesis indicates the number of models tested (see 4.1) and the model names located below the boxplots indicate the models reaching the lowest minimum FDE over the test set. The dashed red line at y = 0 materializes the score of the best performing baseline.

We also reported the number of models tested. Surprisingly, while during the training step univariate input models reached lower MSE than multivariate inputs models for longer time horizons, during the testing phase, for all time horizons, the lowest FDE model is a multivariate model, and visual inspection of the boxplots indicates multivariate models outperform univariate models. These observations suggest univariate models over longer time horizons are likely overfitting. Due to the lower number of actions and sequences used in longer time horizons (see Table 2), we are probably missing data to validate findings over longer periods.

In line with what we observed during the training step, Crossformer (Zhang & Yan, 2023) models achieve the best performance in shorter time horizon in both univariate and multivariate inputs, while in longer time horizons, no model reached the same dominance. The good performance of Crossformer (Zhang & Yan, 2023) indicates the relevance of using a cross-dimension attention mechanism: the consideration of interactions between the players increases the ability to model and predict performance.

## Conclusion

This study presents several novel contributions to the field of sports analytics. First, we leveraged existing work in time series analysis to model short-term performance in Rugby Union. Second, we constructed a dataset using a preprocessing pipeline that includes a method to estimate ball position, addressing a common limitation in sports analysis. By making our dataset publicly available, we contribute to the advancement of sports analytics research. Furthermore, we propose a comprehensive benchmark of state-of-the-art time series models on this prediction task, highlighting which models are best fitted to predict short-term performance in Rugby Union from tracking data.

On our experiments, we found that multivariate to univariate prediction task outperforms univariate to univariate and baselines, indicating the relevance of using players tracking data to predict short term performance. We used here the position of the ball over the x axis of the field as performance indicator, due to its relevance (see introduction section). This approach could be extended to alternative indicators that capture team dynamics more broadly such as the team’s barycenter, or dynamics of specific subgroups, like the barycenter of the forward pack for Rugby Union (players 1 to 8).

Our work also emphasizes models designed to deal with long sequences and decomposition challenges (e.g. electric consumption and weather forecast) can work on shorter time-periods, with data displaying different characteristics.

Our work presents room for improvement. The robustness of disclosed results might be strengthened, as we only gathered data from a single team and miss data for longer time horizons. Moreover, considering only the offensive team is a clear limitation for two reasons. First, the attacking and defensive teams constantly adapt to one another, we are therefore missing precious information about play development. Second, a team in offense in Rugby Union can either try to advance with (by carrying and / or passing) or without (by kicking) the ball. Only considering actions where the analyzed team is in possession of the ball leads us to fail to capture the success of a team trying to advance without the ball, as sometimes, losing the ball is a desired outcome (e.g. when kicking). Finally, we would benefit from using ball tracking data instead of our ball extraction pipeline, as it would (i) be more precise and (ii) allow us to consider defensive actions.

This study opens up several promising research directions. Performance estimation could support the analysis of interpersonal coordination, including fluctuations in running correlations between players, which have been shown to be an important parameter for success in Rugby Union (Rodrigues et al., 2013). Furthermore, predicted offense outcomes could be used in game simulation and invented scenario to test alternative offensive plays. Our research also relates with multiagent trajectory prediction, which could make us benefit from deep learning architectures specifically designed for spatio-temporal data (Yu et al., 2020, Wu et al., 2023). These models leverage Graph Neural Networks to handle the spatial component of the data and are notably used to model pedestrian trajectories. Additionally, applying explainability techniques,which aim to identify the key factors driving model’s prediction, or generating counterfactuals could provide valuable insights for coaching staffs (Odet et al., 2025). While it has not yet been systematically demonstrated, it is plausible that players contribute unequally to the short-term prediction of ball trajectory. Certain specific players, such as the ball carrier and the supporting runners, are likely to exert a stronger influence on where the ball will go next. Focusing on these players could improve prediction accuracy and model interpretability. For example by modifying the feature vector of a given subset of players (for instance, the forward players during a scrum or a lineout and analyzing the resulting changes in performance prediction, one could gain actionable insights into the quality of the initial movements of these key players. This could be helpful in play development simulations to enhance tactical understanding of players and staffs. Best performing models can also be used as reward models in Reinforcement Learning tasks to identify which collective choreography yields the best outcome.

## Acknowledgments

We thank the US Colomiers Rugby Union club for providing us with the data used in this study. This work was granted access to the HPC resources of CALMIP supercomputing center under the allocation P25003.

